# Estimating Genetic Kin Relationships in Prehistoric Populations

**DOI:** 10.1101/100297

**Authors:** Jose Manuel Monroy Kuhn, Mattias Jakobsson, Torsten Günther

**Affiliations:** Uppsala University, Evolutionary Biology Centre, Department of Organismal Biology and SciLifeLab, Norbyvägen 18C, SE-752 36 Uppsala, Sweden; University of Freiburg, Institute of Biology I (Zoology), Hauptstrasse 1, D-79104 Freiburg, Germany

## Abstract

Archaeogenomic research has proven to be a valuable tool to trace migrations of historic and prehistoric individuals and groups, whereas relationships within a group or burial site have not been investigated to a large extent. Knowing the genetic kinship of historic and prehistoric individuals would give important insights into social structures of ancient and historic cultures. Most archaeogenetic research concerning kinship has been restricted to uniparental markers, while studies using genome-wide information were mainly focused on comparisons between populations. Applications which infer the degree of relationship based on modern-day DNA information typically require diploid genotype data. Low concentration of endogenous DNA, fragmentation and other post-mortem damage to ancient DNA (aDNA) makes the application of such tools unfeasible for most archaeological samples. To infer family relationships for degraded samples, we developed the software READ (Relationship Estimation from Ancient DNA). We show that our heuristic approach can successfully infer up to second degree relationships with as little as 0.1x shotgun coverage per genome for pairs of individuals. We uncover previously unknown relationships among prehistoric individuals by applying READ to published aDNA data from several human remains excavated from different cultural contexts. In particular, we find a group of five closely related males from the same Corded Ware culture site in modern-day Germany, suggesting patrilocality, which highlights the possibility to uncover social structures of ancient populations by applying READ to genome-wide aDNA data.

## Introduction

An individual’s genome is a mosaic of different segments inherited from our various direct ancestors. These segments, shared between individuals, can be referred to as identical by descent (IBD). Knowledge about IBD segments has been used for haplotype phasing [1, 2], heritability estimation [3, 4], population history [5], inference of natural selection [6] and to estimate the degree of biological relationship among individuals [7]. A number of methods have been developed to estimate the degree of biological relationship by inferring IBD from SNP genotype or whole genome sequencing data. The methods for estimating relationship levels implemented in PLINK [8], SNPduo [9], ERSA [10, 11], KING [12], REAP [13] and GRAB [14] greatly benefit from genome wide diploid data, information about phase, recombination maps and population allele frequency, and are sometimes able to successfully infer relationships up to 11th degree [11].

Knowing whether a pair of individuals is directly related or not, and estimating the degree of relationship is of interest in various fields: Genome-wide association studies and population genetic analyses often try to exclude related individuals since they do not represent statistically independent samples; in forensics, archaeology and genealogy, individuals and their relatives can be identified based on DNA extracted from human remains [15, 16]; Breeders and conservation biologists are interested in the relatedness of mating individuals [17, 18]. Current methods present significant limitations for the analysis of degraded samples as they rely on diploid genotype calls, low proportions of missing data and sometimes even phase information. Especially in the fields of forensics and archaeology, postmortem damage results in incomplete data due to low concentrations and fragmentation of endogenous DNA in the sample [19–21]. In archaeology, the analysis of IBD has the potential to provide an independent means to test kinship behavior and social organization [22], but current methods would be restricted to exceptionally well-preserved samples. In forensic science and practice, the dominant approach has been to type several short tandem repeat (STR) markers, which in most cases provide sufficient information for relatedness assessment, but the STRs might be hard to type in degraded samples [23]. In addition to nuclear STRs, mitochondrial and Y-chromosome haplogroups have been widely used to infer family relationships (e.g. [15,16, 24, 25]), although they can only exclude certain direct relationships since most mitochondrial and Y-chromosome haplogroups are relatively common among unrelated individuals. These uniparental markers can be typed from degraded samples, and can be used to exclude maternal or paternal relationships, but not to infer the actual degree of relationship. Genome-wide data, however, can be obtained from degraded samples at a higher success rate than STRs and it can be used to confidently identify individuals [26].

Single Nucleotide Polymorphism (SNP) data can be obtained from genotyping experiments (e.g. SNP arrays or RAD sequencing), targeted capture [27], and whole-genome shotgun sequencing (e.g. [28, 29]). The field of ancient DNA has developed rapidly over the last few years and allowed pivotal studies of the population history of Europe [27–37] and the peopling of the Americas [36, 38, 39]. However, both whole-genome shotgun sequencing (e.g. [29, 31, 32]) and genome-wide SNP capture (e.g. [27, 33]) usually achieve coverages <1x per informative site for most individuals which makes diploid genotype calls at all sites virtually impossible. Methods to infer relationships, however, rely on such ideal data to identify IBD blocks which is a major limitation for applying these methods to ancient DNA data.

However, even low coverage data contain information about the degree of relationship. To utilize this information, we developed READ (Relationship Estimation from Ancient DNA), a heuristic method to infer family relationships up to second degree from samples with extremely low coverage. The method is tested on publicly available data with known relationship, which we sub-sample to resemble the properties of degraded samples. We also apply our method to a number of ancient samples from the literature and confidently classify individual pairs as being related.

## Results

### Method Outline

The input for READ are a set of TPED/TFAM files [8] containing genotype calls for a population. The biallelic SNP sites in that file would usually be from some externally ascertained SNP panel (e.g. Human Origins array or 1000 genomes) and all SNPs are assumed for be pseudo-haploid (i.e. one randomly sampled allele per individual and site) as the low coverage in aDNA studies normally does not allow to call heterozygous genotypes. We then divide the genome into non-overlapping windows of 1 Mbps each and for each pair of individuals calculate the proportion of non-matching alleles inside each window *P*0. Similar to [40, 41], the genome-wide distribution of *P*0 is then normalized using the average *P*0 of an unrelated pair of individuals which accounts for effects of SNP ascertainment and population diversity. Depending on the normalized proportion of shared alleles, each pair of individuals is classified as unrelated, second-degree (i.e. nephew/niece-uncle/aunt, grandparent-grandchild or half-siblings), first-degree (parent-offspring or siblings) or identical individuals/identical twins (Fig. 1). As a method with the goal to classify pairs of individuals, READ always outputs the best fitting degree of relationship. To assess the certainty of each categorization, the distance to the classification cutoffs are expressed as multiples of the standard error of the mean (*Z*).

**Fig 1.**
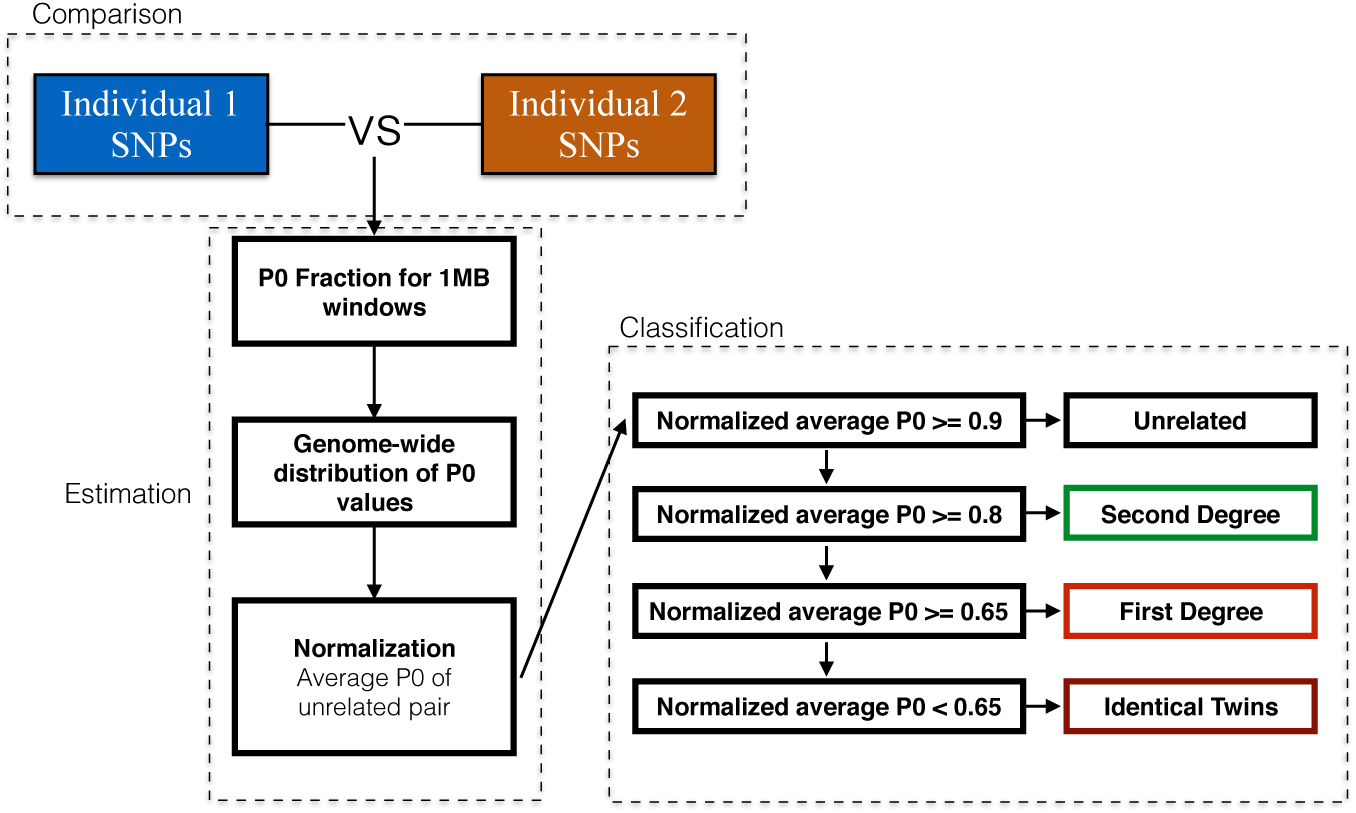
Outline of the general READ workflow to estimate the degree of relationship between two individuals.

### Simulations based on modern data with known relationship

READ’s performance was tested on 1,326 individuals of 15 different populations from the phase 3 data of the 1000 genomes project [42]. A total of 86,336 pairwise comparisons were tested. The rates of false positives (unrelated individuals classified as related) and false negatives (related individuals classified as unrelated) are highly dependent on the amount of data available for pairwise comparison. READ showed an overall good performance with false negative and false positive rates below four percent for as little as 1,000 overlapping SNPs (Fig. 2A). Fig. 3 shows how these SNP numbers would relate to sequencing coverages of the two individuals compared. The proportion of related individuals that were classified as related but not to the correct degree (”Wrong degree”) increased with lower numbers of overlapping SNPs. Separating the error rates between first and second degree relatives shows that most of this increase is due to first degree relatives classified as second degree relatives when the number of SNPs is low (Fig. 2B). False positive rates are low for both degrees of relationship and false negative rate is below one percent for first degree relatives (Fig. 2B and C). The rate of false negatives is considerably high for second degree relatives and it increases up to 39% for low numbers of SNPs (Fig. 2C).

**Fig 2.**
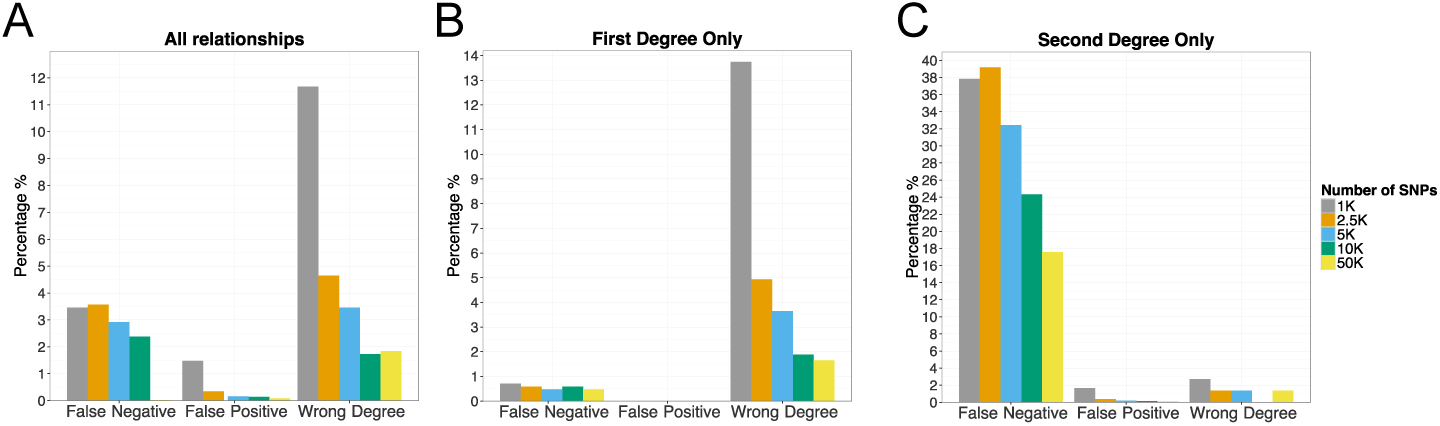
Simulation based estimates of error rates for different numbers of SNPs. The analysis is based on pairs of individuals with known degree of relationship in the 1000 genomes data. (A) All degrees of relationship, (B) only first degree relatives and (C) only second degree relatives. Pairs known to be related which are classified as the wrong degree are shown as”Wrong degree” (e.g. a pair of first degree relatives is classified as second degree relatives).

**Fig 3.**
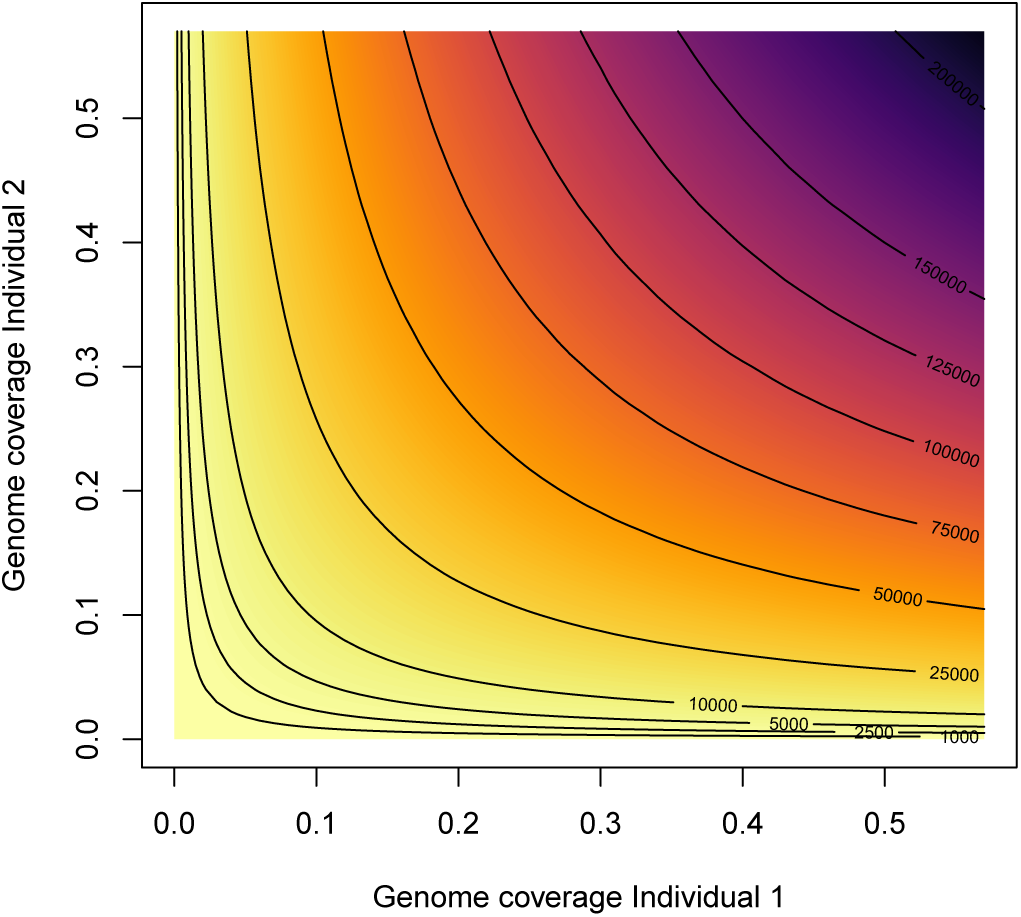
Number of overlapping SNPs dependent on the sequencing coverage for each individual. This figure shows expected number of overlapping SNPs between two individuals for different combinations of sequencing depths. The contour lines mark different numbers of SNPs including those used in the simulations (see Fig. 2). The maximum number of SNPs is set to 1,156,468, identical to what has been used in the simulations and similar to the 1.2 million SNPs used in the empirical data set [33]. The calculation assumes a Poisson distribution of sequencing coverage across the genome [43].

Further complications in the analysis of empirical aDNA data are sequencing and mapping errors, contamination and post-mortem damage. Ultimately, these issues will increase the proportion of wrongly called alleles at SNP sites. To see the effect of such genotyping errors, we repeated the simulations with certain error rates meaning that alleles were randomly changed with a probability corresponding to the defined genotyping error rate. The results of this simulation are shown in Fig. 4. Essentially, wrongly called alleles lead to an overestimation of genetic distance between individuals. As a consequence, pairs of individuals tend to get classified into more distant categories which can be seen by an increase in the proportion of pairs classified as wrong degree and an extremely high false negative rate for higher rates of genotyping error. False positive rates are not affected by wrongly called alleles. Genotype error rates ≤ 5% still seem to produce acceptable false negative rates showing how important it is to keep such errors low in empirical studies. Illumina sequencing has error rates of less than 1% [44–46] and careful data curation as well as filtering (see Discussion) should be able to minimize the impact of other sources of genotyping errors.

**Fig 4.**
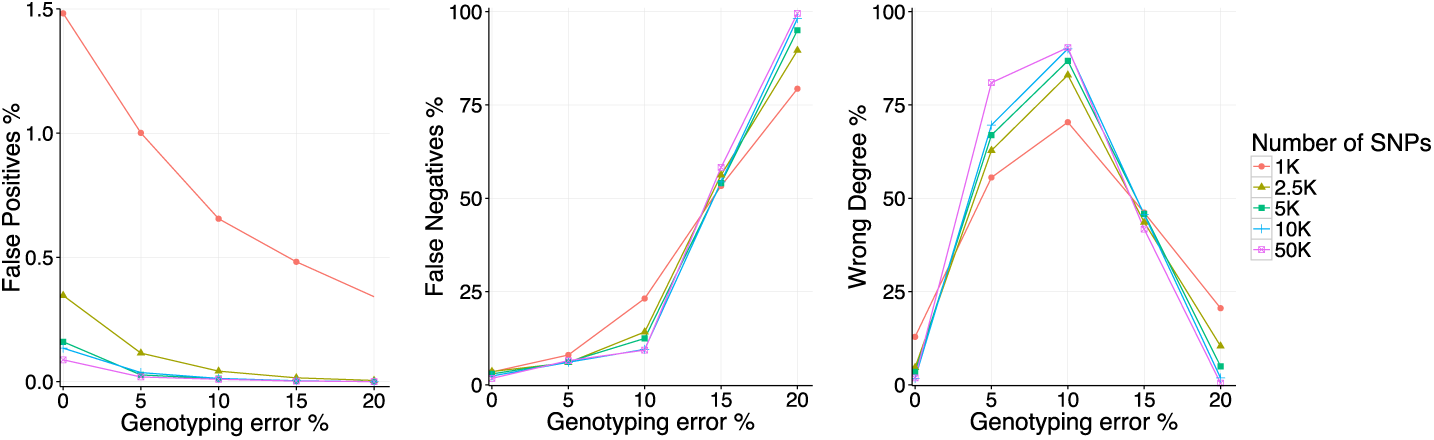
Effect of genotyping error on READ’s performance. The simulations are identical to those conducted for Fig. 2 but including a certain proportion of wrongly called genotypes. The rates of false positives, false negatives and”Wrong degree” were calculated accordingly.

### Relationships among prehistoric Eurasians

To investigate READ’s performance on empirical aDNA data, we analyzed a large published genotype data set of 230 ancient Eurasians from the Mesolithic, Neolithic and Bronze Age periods [33]. In accordance with the original publications [27, 29, 33], READ inferred RISE507 and RISE508 to be the same individual and all nine known relationships were correctly identified as first degree relatives (Table 1). In addition to those, READ identified one additional pair of first degree relatives as well as six new second degree relationships. All relatives are from the same location and their radiocarbon dates (if available) are overlapping.

**Table 1.** Pairs of relatives among the 230 individuals in the aDNA dataset as inferred by READ.

| Group | Ind1 | MT and Y<br>(Ind1) | C14 date<br>(Ind1) | Ind2 | C14 date<br>(Ind2) | MT and Y<br>(Ind2) | Inferred re-<br>lationship | $\vert Z\vert ^{\S}$ |
| --- | --- | --- | --- | --- | --- | --- | --- | --- |
| Afanasievo<sup>S</sup> | RISE507<br>(female) | U5a1a1 | 3322-2923<br>calBCE | RISE508<br>(female) | 3331-2935<br>calBCE | U5a1a1 | identical | 17.16 |
| Neolithic<br>Anatolia<sup>C</sup> | I0736<br>(female) | N1a1a1a | 6500-6200<br>BCE | I0854<br>(female) | 6500-6200<br>BCE | N1a1a1a | 1st | 2.31 |
| Neolithic<br>Anatolia<sup>C</sup> | I1097<br>(male) | W1;<br>G2a2b2a | 6500-6200<br>BCE | I0744<br>(male) | 6500-6200<br>BCE | J1c11;<br>G2a2b2a | 2nd* | 3.50 |
| Bell<br>Beaker,<br>Germany<sup>S</sup> | RISE563<br>(male) | K1c1;<br>R1b1a2a1a2b | NA | RISE564<br>(male) | NA | H; R1b1a2a1 | 2nd* | 0.02 |
| Bell<br>Beaker,<br>Germany<sup>C</sup> | I0111<br>(female) | H3ao | 2475-2204<br>calBCE | I1530<br>(male) | 2345-2198<br>calBCE | H3ao; R1 | 1st | 4.88 |
| Corded<br>Ware,<br>Germany<sup>C</sup> | I1538<br>(male) | J1c5; R1a | 2500-2050<br>BCE | I1534<br>(male) | 2500-2050<br>BCE | K1a1b2a;<br>R(xR1b) | 2nd* | 0.23 |
| Corded<br>Ware,<br>Germany<sup>C</sup> | I1538<br>(male) | J1c5; R1a | 2500-2050<br>BCE | I1541<br>(male) | 2500-2050<br>BCE | U2e1a1; R1a | 1st | 5.51 |
| Corded<br>Ware,<br>Germany<sup>C</sup> | I1539<br>(female) | J1c1b1a | 2625-2291<br>calBCE | I1532<br>(male) | 2500-2050<br>BCE | J1c2e; R1a1a | 2nd* | 1.08 |
| Corded<br>Ware,<br>Germany<sup>C</sup> | I1534<br>(male) | K1a1b2a;<br>R(xR1b) | 2500-2050<br>BCE | I1541<br>(male) | 2500-2050<br>BCE | U2e1a1; R1a | 2nd* | 4.72 |
| Corded<br>Ware,<br>Germany<sup>C</sup> | I1540<br>(male) | J1c5; R1a1 | 2500-2050<br>BCE | I1541<br>(male) | 2500-2050<br>BCE | U2e1a1; R1a | 1st | 5.42 |
| Corded<br>Ware,<br>Germany<sup>C</sup> | I1541<br>(male) | U2e1a1; R1a | 2500-2050<br>BCE | I0104<br>(male) | 2559-2296<br>calBCE | U4b1a1a1;<br>R1a1a1 | 2nd* | 7.26 |
| Chalcolithic<br>Iberia<sup>C</sup> | I1302<br>(male) | J2b1a3;<br>G2a2b2b | 2880-2630<br>BCE | I1314<br>(male) | 2880-2630<br>BCE | J2a1a1; G2a | 1st | 3.15 |
| Chalcolithic<br>Iberia<sup>C</sup> | I1274<br>(male) | H;3 I2a2 | 2880-2630<br>BCE | I1277<br>(male) | 2830-2820<br>calBCE | H3; I2a2a | 1st | 6.21 |
| EN Iberia<sup>C</sup> | I0411<br>(male) | K1a2a; F<sup>§</sup> | 5295-5067<br>calBCE | I0410<br>(male) | 5295-5066<br>calBCE | T2c1d or<br>T2c1d2;<br>R1b1 | 1st | 7.03 |
| Srubnaya<sup>C</sup> | I0421<br>(female) | H3g | 1850-1600<br>BCE | I0430<br>(male) | 1850-1600<br>BCE | H3g;<br>R1a1a1b2a2a | 1st | 6.22 |
| Srubnaya<sup>C</sup> | I0354<sup>¶</sup> (fe-<br>male) | U5a1 | 2016-1692<br>calBCE | I0360<br>(male) | 1850-1200<br>BCE | U5a1; R1a1 | 1st* | 2.99 |
| Unetice<sup>C</sup> | I0117<br>(female) | I3a | 2272-2039<br>calBCE | I0114<br>(male) | 2138-1952<br>calBCE | I3a; I2a2 | 1st | 5.47 |

<sup>S</sup> indicates groups that were shotgun sequenced, <sup>C</sup> indicates SNP capture
<sup>§</sup> showing the lower $|Z|$ of both $Z$ scores (one to the upper threshold, one to the lower threshold)
\* newly reported relationship
<sup>§</sup> potentially haplogroup R, not enough data
<sup>¶</sup> excluded as population outlier in [33]

Combining the information obtained from radiocarbon dating, READ, and uniparental haplotypes can help to narrow down the possible form of relationship. For instance, I0111 (female) and I1530 (male) are inferred (using READ) to be first degree relatives, which means they are either full-siblings, mother/son or father/daughter. The shared mitochondrial haplogroup (H3ao) makes father/daughter less likely (but not impossible), and the slightly older radiocarbon date for I0111 (2475-2204 calBCE versus 2345-2198 calBCE [33]) makes mother/son more likely than siblings while not excluding the latter.

READ identified an unknown pair of first degree relationship between two Srubnaya individuals (I0360 and I0354). Notably, Mathieson et al (2015) [33] have excluded I0354 since she was an outlier compared to other Srubnaya individuals. The shared mitochondrial haplogroup (U5a1) and the slightly older age of I0354 make her the putative mother of I0360. The classification of I0360 and I0354 as first degree relatives is probably genuine considering that READ has very low false positive rates. If this prediction was a false positive, it would be very likely that they are at least second degree relatives as the fraction of unrelated individuals wrongly classified as first degrees is extremely low (Fig. 2B). Furthermore, a highly distinct genetic background of one of the individuals should rather cause false negatives and not false positives, which increases the likelihood that the two individuals are in fact related. I0354 could have been a recent migrant to the region who produced offspring (I0360) with a local male, which would explain both the relationship between I0354 and I0360 and the genomic dissimilarity between I0354 and other Srubnaya individuals.

Particularly interesting is a group of five related males from the Corded Ware site in Esperstedt, Germany (Table 1, Fig. 5). Mathieson et al (2015) [33] described two first degree relationships between I1540 and I1541 as well as between I1541 and I1538. Notably, READ missed the second degree relationship between I1540 and I1538, which is likely to be a false negative as the false negative rate for second degree relatives is known to be substantial with low amounts of data (Fig. 2C) and the value for that pair (0.91) is only 1.2 standard errors above the threshold for second degree relatives (0.9). Identical radiocarbon dates do not help to indicate a chronological order, but based on their Y-chromosomes (all likely R1a, S1 Table), one can suggest that they represent a paternal line of ancestry. I1540 is classified as R1a1, but the Y-chromosomal marker this call is based on (L120) is missing in individuals I1538 and I1541, so they could all carry the same haplotype. In addition to these three individuals, I1534 is a second degree relative of I1538 and I1541, who was carrier of R(xR1b) but a more detailed classification was not possible due to the low coverage. I0104, who is a second degree relative to I1541, might also carry the same Y-chromosome as I1534, I1538, I1540 and I1541, but that cannot be determined due to low coverage in those individuals. Generally, the data would be consistent with all five individuals carrying the same Y-haplotype as there are no contradicting calls for R1a defining markers (S1 Table), which would suggest paternal relationship among them. In total, 13 Corded Ware individuals from Esperstedt were genotyped, nine of them were males. It is notable that all five related Esperstedt individuals discussed here were males and only one pair of related Corded Ware individuals from Esperstedt involved a female (I1539 and I1532; Table 1).

**Fig 5.**
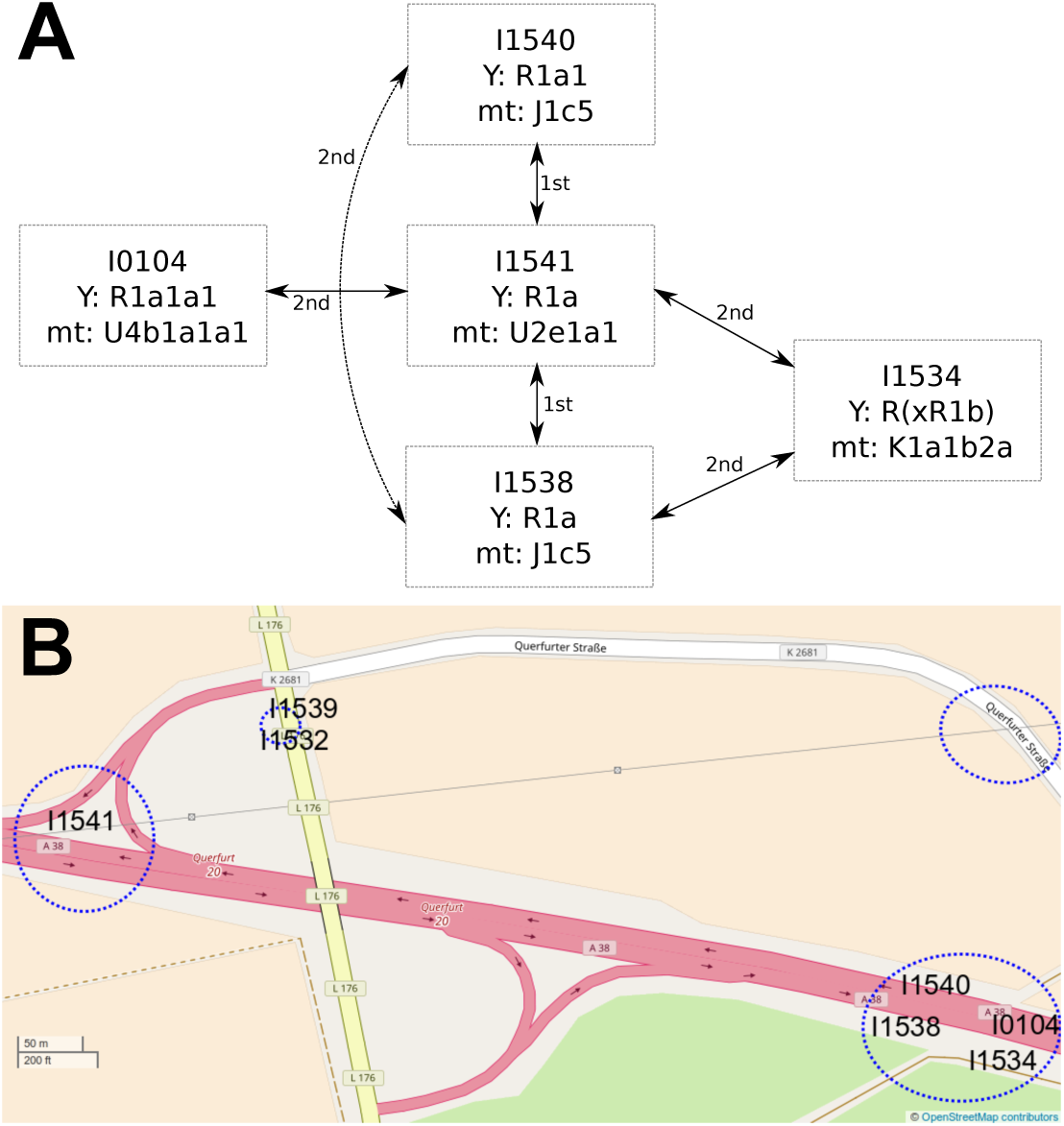
Kin-relationship among males at the Corded Ware site in Esperstedt, Germany. (A) The five individuals, their inferred degree of relationship and their uniparental haplogroups. The dashed line between I1540 and I1538 shows a second degree relationship missed by READ. (B) Map of the Corded Ware site (reference site 4) near Esperstedt, Germany. Blue circles show the locations of Corded Ware Burials. The approximate burial locations of the individuals with inferred relationships are indicated by their ID. Map data © OpenStreetMap contributors, CC BY-SA.

### Normalization in the aDNA data set

READ uses the average *P*0 from an unrelated pair of individuals to normalize the distribution for all test individuals. For our empirical data analysis, we assumed the median of all average *P*0 across pairs of individuals within a test population to represent unrelated individuals, as high values may be caused by recent migrants and low values by related individuals. Fig. 6 shows the distributions of all average P0 before normalization highlighting that the populations exhibit different degrees of background diversity. It is also apparent how the pairs of related individuals (see Tab. 1) are outliers with lower pairwise differences. Most groups from similar geographic and cultural groups show similar medians. These include Neolithic groups (except Iberia EN) and Yamnaya, and – to some degree – Late Neolithic and Bronze Age central Europeans. The latter set of populations could almost belong to two subgroups which cluster by data type (shotgun versus capture) instead of archaeological culture (Unetice, Corded Ware and Bell Beaker). This difference was not observed in Yamnaya for which both data types exist as well. The discrepancy highlights a potential risk of batch effects which has its consequences for the application of READ. Overestimating the distance between unrelated individuals could overestimate relationships in the test group and consequently cause false positives while underestimating the distance between unrelated individuals would have the opposite effect. The extent of the misclassification would be proportional to the ratio between true and used normalization value. For example, if the true value was 0.22 (e.g. Motala HG, Fig. 6) but 0.25 was used (e.g. Hungary EN), an unrelated pair of individuals could be classified as second degree relatives (0.22/0.25 = 0.88 < 0.9). Using the shotgun Bell Beaker median (0.245) to normalize the captured Bell Beaker data does not cause any changes in the classifications, whereas using the capture Bell Beaker median (0.257) for the shotgun data would classify RISE563 and RISE564 as second degree relatives. These two individuals might actually be related, but the value used for normalization would be higher than any pairwise comparison within the shotgun sequenced Bell Beakers. This violates the assumption that the normalization value represents the expectation for a pair of unrelated individuals so this result should be considered a false positive due to a batch effect.

**Fig 6.**
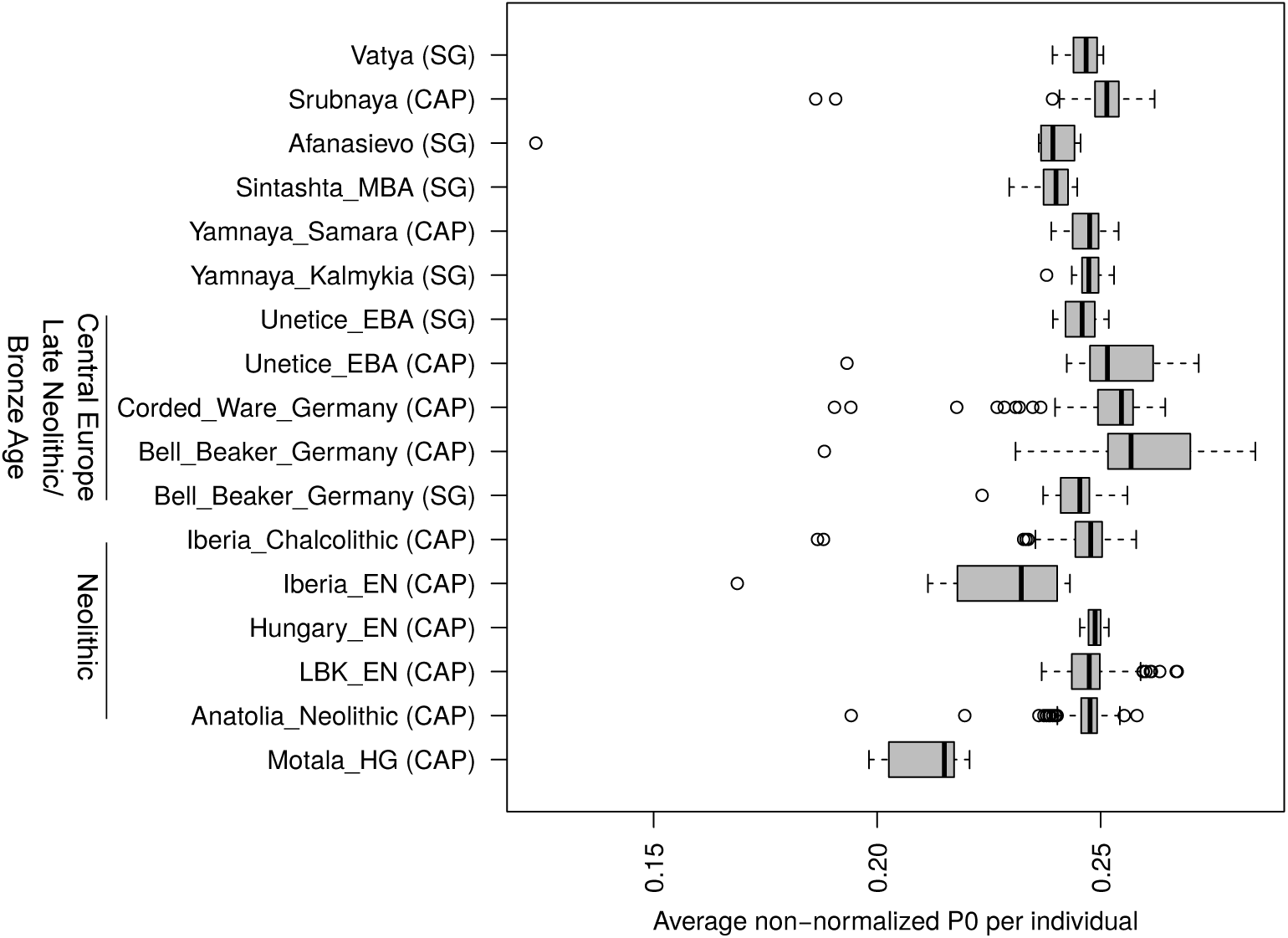
Population distributions of average *P*0 before normalization. The boxplots show all non-normalized average *P*0 scores (one per pair of individuals) per culture. CAP and SG indicate whether the individuals were subject to SNP capture or shotgun sequencing, respectively. A broader chronological/geographical context is shown on the left.

## Discussion

### Applying READ to aDNA data

Several methods to estimate the degree of relationship between pairs of individuals have been developed. For genome-wide diploid data with low error rates, they successfully infer relationships up to 11th degree [11]. Since such data cannot be obtained from degraded samples, a loss in precision was expected. Estimation of second degree relationships (i.e. niece/nephew-aunt/uncle, grandparent-grandchild, half-siblings) is sufficient to identify individuals belonging to a core family which were buried together. We can show that obtaining as little as 2,500 overlapping common SNPs is enough to classify up to second degree relationships from effectively haploid data. The biggest limitations when using such low numbers of SNPs is the high rate of false negatives for second degree relatives. READ can be considered as a conservative tool that avoids false positives by having a relatively high false negative rate which can be decreased substantially with more data. Missing some second degree relationships seems preferable over wrongly inferring relationships for unrelated individuals. A consequent advantage of our method is that it is very unlikely that first degree relatives are classified as unrelated but some second degree relatives might be wrongly classified as unrelated. Shared uniparental haplotypes or a test result close to the threshold (e.g. less than two standard errors difference) could raise such suspicions and might motivate additional sequencing of the samples in question. The amount of overlapping SNPs depends on the genome coverage of both individuals (Fig. 3; e.g. two 0.1x individuals will have approximately the same amount of overlapping data as a 0.05x and a 0.2x individual or a 0.01x individual and a 1x individual). The number of SNPs required for a positive classification as first degree can be obtained by shotgun sequencing all individuals to an average genome coverage of 0.1x (Fig. 3), which is in reach for most archaeological samples displaying some authentic DNA. More data would be beneficial to avoid false negatives in the case of second degree relatives. Recently developed methods for modern DNA, which use genotype-likelihoods to handle the uncertainty of low to medium coverage data require 1-3x genome coverage to estimate third degree relationships [47–49]. Such approaches are promising for well-preserved samples but these coverages might not be within reach for most aDNA studies. Other methods specifically designed for ancient DNA data either require larger population sample sizes than READ [50], large reference data sets [41,51] or are not directly designed to identify relatives and estimate their degrees [52].

READ does not explicitly model aDNA damage and it only considers one allele at heterozygous sites. This implies that a careful curation of the data is required to avoid errors due to low coverage, short sequence fragments, deamination damage, sequencing errors and potential contamination. We recommend a number of well established filtering steps when working with low coverage aDNA data [27–33, 53, 54]. To avoid batch effects, all samples should be processed as similar as possible – at least the bioinformatic pipeline should be identical for all samples. Only fragments of 35 bp or longer should be mapped to the human genome as shorter fragments might represent spuriously mapping microbial contamination [55, 56]. All downstream analysis should be restricted to reads and bases with mapping and base qualities of 30 or higher to reduce the potential effects of mismapping and sequencing errors [56, 57]. To further reduce the effect of sequencing errors, most aDNA studies only consider biallelic SNPs known to be polymorphic in other populations, and call pseudo-haploid genotypes by randomly sampling one read covering that position. Deamination damage can be avoided during the data generation by enzymatic repair of damages [58], or later by computational rescaling of base qualities before SNP calling [59] or by excluding all transition SNPs. For humans, millions of polymorphic transversion sites are known across the genome [42] still leaving substantial amounts of data for analyzing such data sets. Furthermore, a range of methods exist to estimate human contamination of a particular sample [60–64] and the analysis could be restricted to fragments displaying characteristic damage to filter contamination [65, 66]. Finally, each study could simulate data exactly resembling the empirical data analyzed (fragment sizes, damages, contamination) to evaluate how these factors would affect the downstream analysis [56].

An important part of the READ pipeline is the normalization step. This step makes the classification independent of within population diversity, SNP ascertainment and marker density. This property, however, requires at least one additional and unrelated individual from the same population and ideally the same data type to avoid batch effects. The assignment of all individuals to a population can be checked with established methods as principal component analysis (PCA) or outgroup *f*_3_ statistics [39]. Alternatively, a pair of individuals from a different population with similar expected diversity could be used for normalization. Fig. 6 shows that most (but not all) groups from similar cultural and geographical backgrounds have relatively similar normalization scores, but caution should be taken as strong misspecification of the normalization value can cause false negatives or false positives (see Results section). In practice, the relationships are not known a priori. For our data analysis, we assumed that the median across all pairs of individuals from a population of more than four samples represents a proxy of an unrelated pair (as the number of pairs is 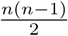; e.g. 10 pairs for a sample size of 5), which we also set as the default mode for READ. The implementation of READ offers other options as well since the median would not work in cases like parent-child-trios (two first degree relationships, one unrelated), where the maximum of all three comparisons should be used for normalization. Other methods normalize by obtaining allele frequency data for a whole population [47, 51], which seems less feasible than obtaining just one unrelated individual (or a pair of unrelated individuals from a surrogate population). Furthermore, prehistoric populations are quite differentiated from modern groups [31, 37, 53] so using modern populations as references for the allele frequencies might introduce biases. A certain limitation for all kinship estimation methods is if the sampled population itself cannot be considered homogeneous, for example due to varying degrees of admixture. Only quite recent developments in inferring relationships can efficiently deal with such cases for modern data [67].

### Kinship in prehistoric populations

We successfully applied READ to data obtained from ancient individuals. READ confidently found all known relationships in the dataset. Furthermore, it identified a number of previously unknown relationships, mainly of second degree. The combination of genomic data, uniparental markers and radiocarbon dating allowed us to infer how two individuals were related to each other. Additional information such as osteological data on the age of the samples or stratigraphic information as burial location or depth could further help to assess the direction of a kinship. Of particular interest was a group of five males from Esperstedt in Germany who were associated with the Corded Ware culture – a culture that arose after large scale migrations of males [68] from the east [27, 29]. Around 50 Corded Ware burials (Fig. 5B), six of them stone cists, were excavated near Esperstedt in the context of road constructions in 2005 [27, 69]. Characteristic Corded Ware pottery was found in the graves and all male individuals had been buried on their right hand site [69]. Interestingly, the central individual of the group of related individuals (I1541, Fig. 5A) was buried in a stone cist approximately 700 meters from the graves of the other four individuals which were all close to each other (Fig. 5B) [69]. The close relationship of this group of only male individuals from the same location suggest patrilocality and female exogamy, a pattern which has also been found from Strontium isotopes at another Corded Ware site just 30 kilometers from Esperstedt [15] and suggested for the Corded Ware culture in general [70]. This represents just one example of how the genetic analysis of relationships can be used to uncover and understand social structures in ancient populations. More data from additional sites, cultures and species other than humans will offer various opportunities for the analysis of relationships based on genome-wide data.

## Materials and Methods

### Approach to detect related individuals

Our approach is based on the methodology used by GRAB [14] which was designed for unphased and diploid genotype or sequencing data. This approach divides the genome into non-overlapping windows of 1 Mbps each and compares for a pair of individuals the alleles inside each window. Each SNP is classified into three different categories: IBS2 when the two alleles are shared, IBS1 when only one allele is shared and IBS0 when no allele is shared. The program calculates the fractions for each category (*P*2, *P*1 and *P*0) per window and, based on certain thresholds, uses them for relationship estimation. GRAB can estimate relationships from 1st to 5th degree, but it has not been tested with data from different SNP panels or populations [14].

We assume that our input data stems from whole genome shotgun sequencing of an ancient sample resulting in low coverage sequencing data. Therefore, we only expect to observe one allele per individual and site which is either shared or not shared between the two individuals. READ does not model aDNA damage, so it is expected that the input is carefully filtered, e.g. by restricting to sites known to be polymorphic, by excluding transition sites or by rescaling base qualities before SNP calling [59]. Analogous to GRAB [14], we partition the genome in non-overlapping windows of 1 Mbps and calculate the proportions of haploid mismatches and matches, *P*0 and *P*1, for each window. Since *P*0 + *P*1 = 1, we can use *P*0 as a single test statistic. The average *P*0 is calculated from the genome-wide distribution. To reduce the effect of SNP ascertainment, population diversity and potential batch effects, each individual pair’s average *P*0 scores are then normalized by dividing all values by the average non-normalized *P*0 score from an unrelated pair of individuals from the same population ascertained in the same way as for the tested pairs. Such a normalization step is not implemented in GRAB [14] suggesting that GRAB might be sensitive to ascertainment bias and general population diversity. The normalization sets the expected score for an unrelated pair to 1 and we can define classification cutoffs which are independent of the diversity within the particular data set. We define three thresholds to identify pairwise relatedness as unrelated, second-degree (i.e. nephew/niece-uncle/aunt, grandparent-grandchild or half-siblings), first-degree (parent-offspring or siblings) and identical individuals/identical twins. The general work flow and the decision tree used to classify relationships is shown in Fig. 1. There are four possible outcomes when running READ: unrelated (normalized P0≥0.9), second degree (0.9≥normalized P0≥0.8), first degree (0.8≥normalized P0≥0.65) and identical twins/identical individuals (normalized P0<0.65) (Fig. 1). The cutoffs were chosen to maximize precision in the pseudo-haploidized 1000 genomes dataset (see below) before randomly subsampling SNPs. These values are similar to the probabilities of one randomly chosen allele for an individual being IBD to a randomly chosen allele from another individual considering their degree of relationship. The option of classifying two individuals as third degree was not implemented as the few known third degree relationships in the empirical datasets showed values similar to unrelated individuals. READ is implemented to classify pairs of individuals in certain categories, so it will always output the best fitting degree of relationship. As a measurement of confidence of that classification, we estimate the standard error of the mean of the distribution of normalized *P*0 scores and calculate the distance to the cutoffs in multiples of the standard error (similar to a *Z* score).

Relationship Estimation from Ancient DNA (READ) was implemented in Python 2.7 [71] and GNU R [72]. The input format is TPED/TFAM [8] and READ is publicly available from https://bitbucket.org/tguenther/read

### Modern data with reported degrees of relationships

Autosomal Illumina Omni2.5M chip genotype calls from 1326 individuals from 15 different populations were obtained from the 1000 genomes project (ftp://ftp.1000genomes.ebi.ac.uk/vol1/ftp/release/20130502/supporting/hd_genotype_chip/) [42]. We used vcftools version 0.1.11 [73] to extract autosomal biallelic SNPs with a minor allele frequency of at least 10% (1,156,468 SNPs in total – similar to the aDNA data set used for the empirical data analysis [33]; see below) and to convert the data to TPED/TFAM files. The data set contains pairs of individuals that were reported as related, 851 of them as first degree relationships and 74 as second degree. We randomly sub-sampled 1000, 2500, 5000 and 50000 SNPs and also randomly picked one allele per site in order to mimic extremely low coverage sequencing of ancient samples. READ was then applied to these reduced data sets and the median of all average *P*0s per population was used to normalize scores assuming that this would represent an unrelated pair. Individual pairs with known relationship and their degree of relatedness are shown in S2 Table and S3 Table. Additionaly, we introduced different error rates to the data to assess the possible effects of sequencing and mapping errors, contamination and post-mortem damage. Error rates were introduced by randomly changing alleles to the alternative with probabilities of 5, 10, 15 and 20%. Related individuals classified by READ as unrelated were considered as false negatives, unrelated individuals classified as related were considered as false positives and related individuals classified as related but not on the proper degree were considered as wrong degree. The false negative rate was obtained by dividing the number of false negatives by the total number of true related pairs, the false positive rate by dividing the number of false positives by the total number of unrelated pairs and the wrong degree rate by dividing the number of incorrectly classified related pairs by the total number of true related pairs.

### Ancient data

In addition to the modern data, published ancient data was obtained from the study of Mathieson et al. (2015) [33]. The data set consisted of 230 ancient Europeans from a number of publications [27, 29–31, 54, 74] as well as new individuals from various time periods during the last 8,500 years. The data set consisted of haploid data for up to 1,209,114 SNPs per individual. We extracted only autosomal data for all individuals and applied READ to each cultural or geographical group (as defined in the original data set of Mathieson et al (2015) [33]) with more than four individuals separately. Shotgun sequencing data was also analyzed separately from SNP capture data to avoid batch effects. The median of all average *P*0s per group was used for normalization assuming that this would represent an unrelated pair. Mathieson et al (2015) [33] report nine pairs of related individuals and they infer all of them to be first degree relatives without providing details on how they were classified. Y-chromosome haplotypes of the five individuals shown in Fig. 5A were checked using samtools [75] (applying a minimum mapping and base quality of 30) and marker information for the haplotypes R1a and R1b from the International Society of Genetic Genealogy (http://www.isogg.org, accessed January 16, 2017). The results are shown in S1 Table.

## Supporting Information

**S1 Table. Y chromosome calls for haplogroup R defining markers in the five individuals shown in Fig. 5A.**

**S2 Table. Pairs of first degree related individuals in the 1000 genomes data.**

**S3 Table. Pairs of second degree related individuals in the 1000 genomes data.**

## Acknowledgments

We thank Federico Sanchez-Quinto, Jan Storá, Rita Peyroteo Stjerna, four anonymous reviewers and those who commented on the preprint for constructive feedback on the manuscript as well as Gülşah Merve Dal Kılınç and Mehmet Somel for discussions on the approach. Computations were performed on resources provided by SNIC through Uppsala Multidisciplinary Center for Advanced Computational Science (UPPMAX) under projects b2013203 and b2015094.

